# HAPNEST: efficient, large-scale generation and evaluation of synthetic datasets for genotypes and phenotypes

**DOI:** 10.1101/2022.12.22.521552

**Authors:** Sophie Wharrie, Zhiyu Yang, Vishnu Raj, Remo Monti, Rahul Gupta, Ying Wang, Alicia Martin, Luke J O’Connor, Samuel Kaski, Pekka Marttinen, Pier Francesco Palamara, Christoph Lippert, Andrea Ganna, Intervene Consortium

## Abstract

Existing methods for simulating synthetic genotype and phenotype datasets have limited scalability, constraining their usability for large-scale analyses. Moreover, a systematic approach for evaluating synthetic data quality and a benchmark synthetic dataset for developing and evaluating methods for polygenic risk scores are lacking. We present HAPNEST, a novel approach for efficiently generating diverse individual-level genotypic and phenotypic data. In comparison to alternative methods, HAPNEST shows faster computational speed and a lower degree of relatedness with reference panels, while generating datasets that preserve key statistical properties of real data. These desirable synthetic data properties enabled us to generate 6.8 million common variants and nine phenotypes with varying degrees of heritability and polygenicity across 1 million individuals. We demonstrate how HAPNEST can facilitate biobank-scale analyses through the comparison of seven methods to generate polygenic risk scoring across multiple ancestry groups and different genetic architectures.

## 1 Introduction

With the emergence of large-scale biobanks, methods to analyse common genetic variants (single nucleotide polymorphisms, or SNPs) across diverse human populations are in growing demand. This is especially the case for polygenic risk scoring (PRS) methods, which quantify an individual’s genetic risk for a disease or other phenotypic trait [1]. Derived from one’s genotype, well-calibrated PRSs have the potential to be used for risk stratification and prognostic prediction [1]. PRS’s utility has been demonstrated for certain common diseases among European ancestries, on which most genome-wide association studies (GWAS) were carried out [2], but some studies have highlighted limitations in PRS’s transferability across ancestries and different socio-demographic groups [3]. Thus, the development of methods that can improve the generalisability of PRSs is needed. At the same time, only a few accessible large-scale biobank datasets exist and most previous PRS methods have been tested and compared in UK Biobank [4]. More diverse biobank datasets are needed, but due to the highly sensitive nature of genetics data, accessing and sharing individual-level data raises privacy concerns. This makes publicly accessible synthetic data a welcome alternative for methods developers.

Broadly, two main approaches have been used to simulate individual level genetic data. Coalescence based methods, such as Hudson’s ms and msprime [5, 6], use demographic models to generate genomes including both rare and common variants. Reference-based approaches use real genomic data (e.g. 1000 genomes or HGDP) to generate synthetic data, but they are not suitable to generate realistic rare variants. There are also methods, such as simGWAS[7], that directly simulate GWAS summary statistics. However, many times they do not meet modern demands for methods development based on individual level data. We will focus on reference-based approaches since for PRSs we are mostly interested in common genetic variation, which forms the bulk of complex trait heritability [8]. Moreover, common SNPs, especially Hapmap3 SNPs [9], are widely recommended for PRS computation [10]. HAPGEN2 [11] is a widely used tool for genotype and phenotype simulation, which preserves linkage disequilibrium (LD) patterns of real data through a resampling approach based on the Li and Stephens model [12]. However, HAPGEN2 lacks computational scalability and flexibility to simulate certain scenarios of interest for biobank-scale PRS and SNP-based methods development. Recent alternatives include G2P [13] and Sim1000G [14]. Sim1000G is an integrated R package, but is limited to genotype simulation. G2P encompasses both genotype and phenotype simulation, and is highly customisable, but this setup can be challenging for non-expert users. Without an integrated approach for parameter selection and evaluation of synthetic data quality, it is difficult for end-users to understand the statistical guarantees and reliability of the generated datasets. To the best of our knowledge, there does not exist a software tool implementing an end-to-end pipeline for synthetic data generation, evaluation and optimisation.

To address these limitations, we introduce HAPNEST, a user-friendly tool for generating synthetic datasets for genotypes and phenotypes, evaluating synthetic data quality, and analysing the behavior of model parameters with respect to the evaluation metrics. HAPNEST simulates genotypes by resampling a set of existing reference genomes, according to a stochastic model that approximates the underlying processes of coalescent, recombination and mutation. It is, in spirit, similar to HAPGEN2, but we introduce some innovations to reduce relatedness between synthetic individuals and the reference panel. Phenotypes are subsequently assigned to each sample by integrating user-specified genetic, covariate, and environmental effects. Genetic effects are modelled in terms of heritability and polygenicity. HAPNEST enables efficient simulation of diverse biobank-scale datasets, as well as simultaneously generating multiple genetically correlated traits with population specific effects under different pleiotropy models. Moreover, the HAPNEST software includes an extensive workflow for evaluating synthetic data fidelity and generalisability, as well as approximate Bayesian computation (ABC) techniques for analysing the posterior distributions of model parameters to aid model selection.

We compare the performance of HAPNEST with current state-of-the-art genotype and phenotype simulation tools in terms of data quality and computational speed. Furthermore, as a demonstration of the utility of our tool, we show the application of our diverse, biobank-scale synthetic data for evaluating the performance of various PRS methods under different disease models. Our open source software tool is available at https://github.com/intervene-EU-H2020/synthetic_data, and has also been distributed as Docker and Singularity containers. We have generated 6.8 million common variants and 9 phenotypes with varying degrees of heritability and polygenicity across 1 million individuals and made this large synthetic dataset available at https://www.ebi.ac.uk/biostudies/studies/S-BSST936 to encourage standardised evaluation of new statistical methods by the genomic research community.

## 2 Results

### 2.1 Overview of genotype generation methods

Synthetic haplotypes are constructed as a mosaic of segments of various lengths imperfectly copied from real haplotypes (Figure 1, Panel a). HAPNEST uses an approximate model inspired by the sequential Markovian coalescent model [15], which makes simplifying assumptions about the coalescence and recombination processes. The real haplotypes to copy from are sampled uniformly from a reference dataset, 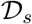, limited to individuals belonging to a certain ancestry group *s*. Alternatively, users can specify the proportion of real haplotypes to sample from each ancestry group. We refer to the Discussion section of the paper regarding the complications in interpreting admixed samples. Segments of length *ℓ* (in centimorgans) are sampled from the real haplotypes (Figure 1, Panel b) based on a simplified stochastic model of the coalescent and recombination processes,

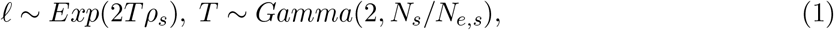

where *ρ_s_* is the population-specific recombination rate, *N_e,s_* is the population-specific mean effective population size, and *N_s_* is the number of reference samples for population *s*. The simulation of varying, rather than constant, coalescence time *T*, is one of two main aspects in which HAPNEST differs from previous methods such as HAPGEN2. Another feature we introduce is that to reduce close copying of genotypes from the reference, the presence of a genetic variant at position *i* is only copied if *T* ≤ *m_i_*, where *m_i_* is the variant’s age of mutation (obtained from [16]). Two synthetic haplotypes, *h_i_*, *i* ∈ {1, 2}, constructed in this way are added element-wise to create a synthetic genotype, *g* (Figure 1, Panel c). For experiments in this text, we consider a reference dataset of 4,062 phased genotypes derived from the publicly available 1,000 Genomes Project and Human Genome Diversity Project datasets for 6 major discrete ancestry groups [17].

**Figure 1:**
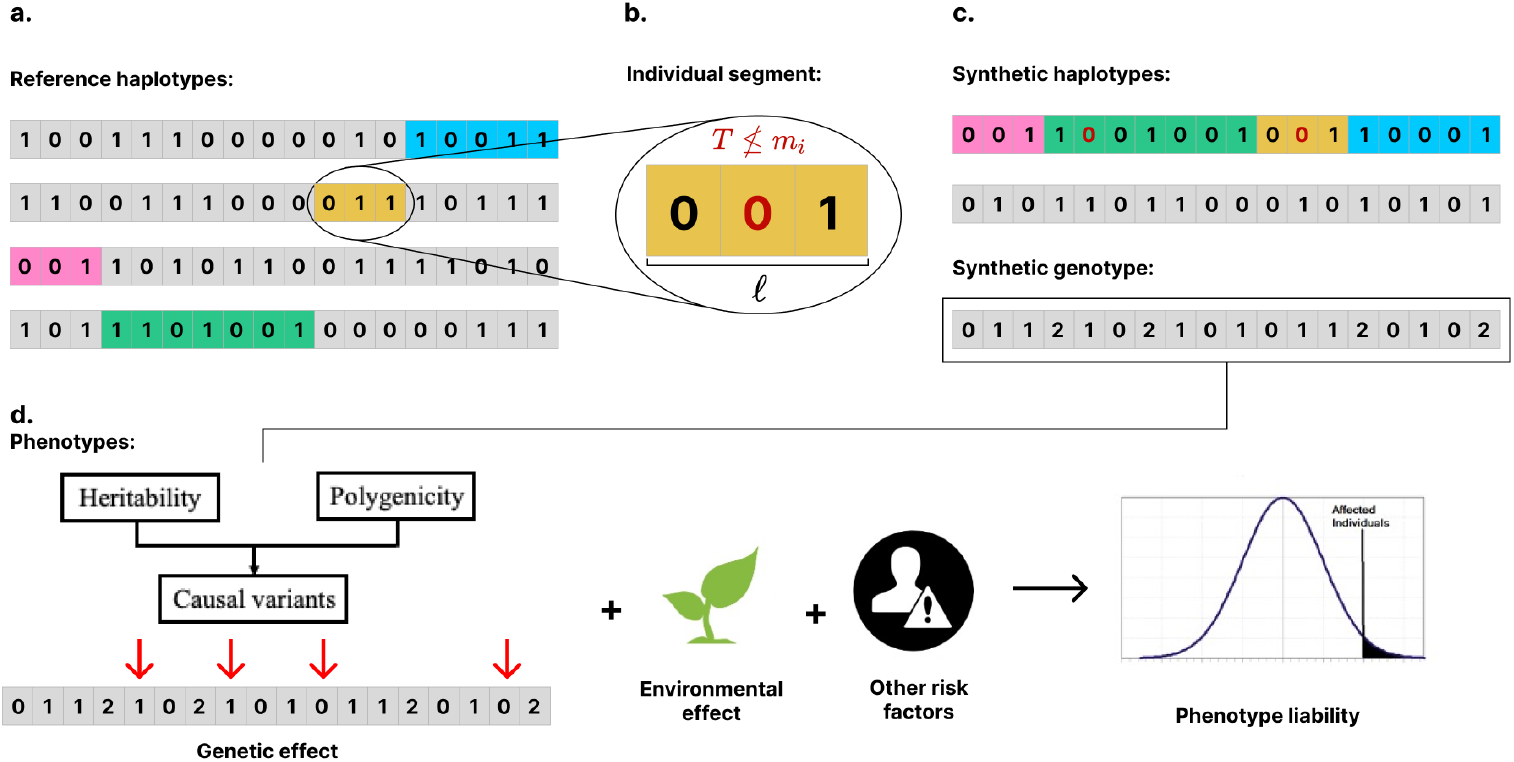
**a**. A reference set of real haplotypes, from which segments (colored) are imperfectly copied to construct a synthetic haplotype. **b**. Detailed view of an individual segment. The segment length, *ℓ*, and coalescence time, *T*, are sampled from a stochastic model. The presence of a genetic variant at position *i* is only copied if *T* ≤ *m_i_*, where *m_i_* is the variant’s age of mutation. Variants that are not copied are shown in red. **c**. Synthetic genotypes, *g*, are constructed as pairs of synthetic haplotypes, *h_i_*, *i* ∈ {1, 2}. **d**. Once the genotype is generated, liability of phenotype will subsequently be assigned to each sample as a summation of genetic effect, covariate effect (if any) and environmental noise.

### 2.2 Posterior distributions of model parameters

While there is no consensus on universal metrics for evaluating synthetic datasets, the literature tends to emphasise the general properties of fidelity (the ability to preserve statistical properties of the real data) and generalisability (the extent to which synthetic samples are not direct copies of the real data) [18, 19]. For downstream applications such as GWAS and PRS, it is important to preserve realistic LD patterns in the synthetic data (fidelity objective). In this work we measure generalisability in terms of genetic relatedness (defined by the kinship coefficient), to ensure that the samples in large synthetic datasets are not close copies of samples from the much smaller reference dataset. We use Approximate Bayesian Computation (ABC) (as explained in the Methods section) to estimate model parameters which balance both the fidelity and the generalisability objectives. Figure 2 shows the posterior distributions of the parameters that best satisfy these criteria. We observe a tradeoff between optimising the fidelity objective (Supplementary, Figure 10) and optimising the generalisability objective (Supplementary, Figure 11). This tradeoff can affect the results of downstream analyses such as GWAS (Supplementary, figure 12, table 6).

**Figure 2:**
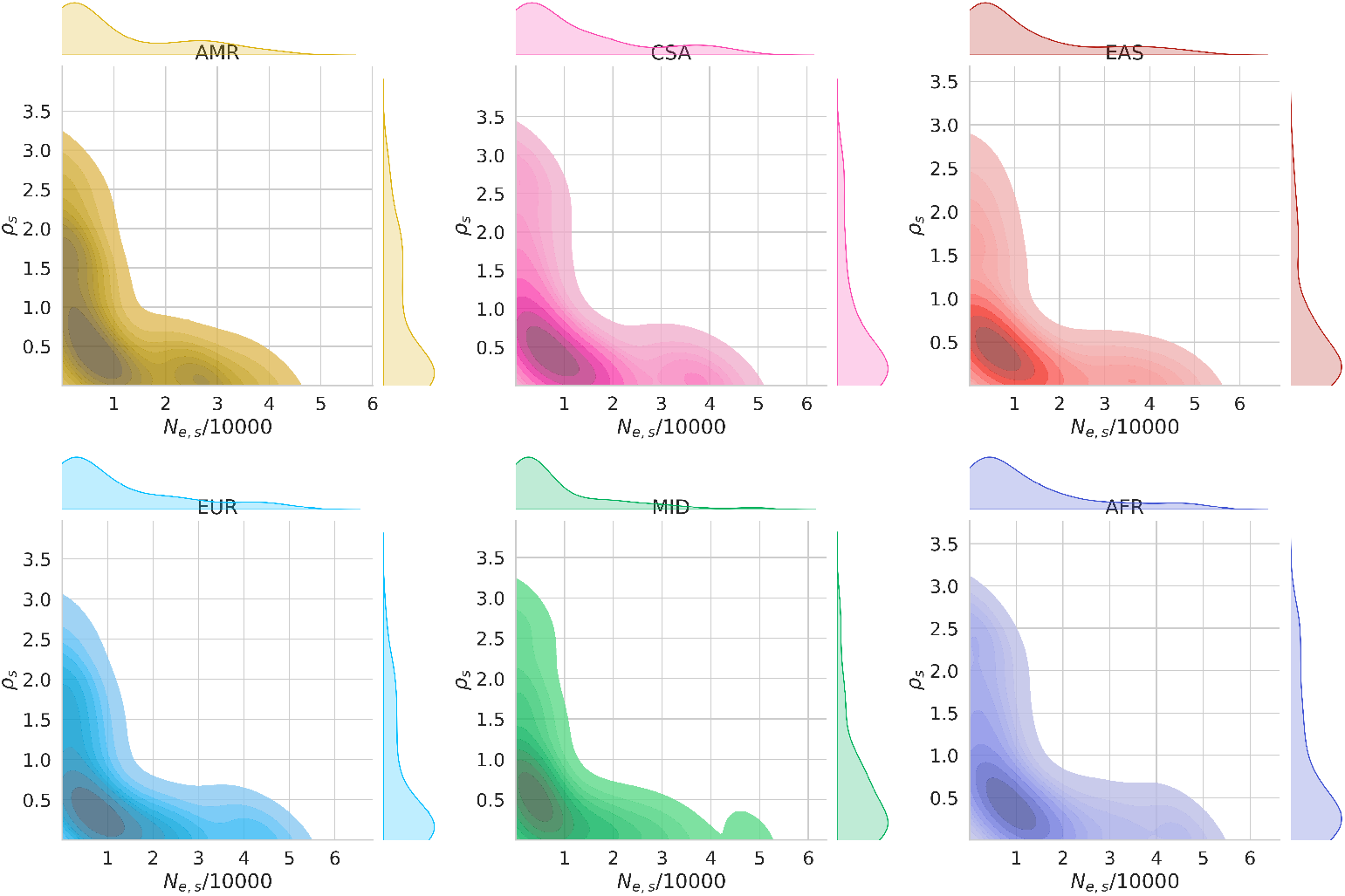
Posterior distributions plotted as marginal and bivariate kernel density estimates for the effective population size, *N_e,s_*, and recombination rate, *ρ_s_*, for six superpopulation groups, *s* ^1^. The experiment setup used 500 simulations for *N_syn_* = 1000 synthetic samples based on a reference of chromosome 21 HapMap3 variants, with uniform priors and a 20 percent rejection rate.

### 2.3 Comparison of synthetic genotype quality

Synthetic data quality is evaluated based on a workflow implemented in the HAPNEST software tool for measuring the fidelity, diversity and generalisability of synthetic datasets. Briefly, fidelity is measured as the similarity between the real (reference) and synthetic datasets for 4 properties: minor allele frequency (MAF) distribution, population structure in terms of alignment of the principal components (PCs), LD decay and nearest neighbour adversarial accuracy (as explained in the Methods section). Diversity is measured by the degree of genetic relatedness (kinship) within the synthetic dataset and generalisability is measured by the degree of genetic relatedness between the real and synthetic datasets. HAPNEST is compared with three alternative methods (HAPGEN2, G2P and Sim1000G) for *N_syn_* = 1, 000 synthetic samples, based on a reference dataset of *N_ref_* = 775 European-ancestry individuals. In this section we compare two parameter sets for HAPNEST: HAPNEST-abc, as determined by the ABC procedure for balancing the LD and relatedness objectives (Figure 2) and HAPNEST-equivalent, that is more equivalent to the parameter configurations used by the other tools (which do not have built-in optimisation procedures). The rest of the text generally considers the ABC parameters, unless otherwise stated.

#### 2.3.1 Fidelity

The full fidelity results are reported in Supplementary Table 1. The HAPNEST-equivalent and G2P methods had the lowest divergence in MAF between the synthetic and reference datasets, followed by HAPNEST-abc, Sim1000G and HAPGEN2. The HAPNEST-equivalent method also had the LD decay that least diverged from the reference, followed by the G2P and HAPGEN2 methods. However, we observe that HAPNEST-abc has a faster LD decay (Figure 3, Panel b) and more generally, our posterior analysis indicates there is a tradeoff between optimizing the LD and relatedness objectives (Supplementary, Figure 10, 11). Nevertheless, GWAS results presented later still indicate realistic LD structure at genome-wide significant loci. We evaluate preservation of population structure by comparing the PC alignment score, defined as the cosine distance between the first 20 PCs obtained from real and synthetic data within European individuals. HAPGEN2 has the highest PC alignment score, followed by HAPNEST-equivalent. HAPNEST can also generate datasets that preserve population structure across multiple populations (Figure 4, Panel a). Finally, we consider privacy-preserving metrics, by calculating the nearest neighbour adversarial accuracy score, which averages the true positive rate and true negative rate for distinguishing real and synthetic data. Adversarial accuracy scores closest to 0.5 are observed for the G2P and HAPNEST-abc methods, indicating that these synthetic samples are more indistinguishable from the real data. Our analysis indicates that no one method performs best across all evaluation metrics, but instead there are tradeoffs that end users should consider, depending on the priorities of their use case.

**Figure 3:**
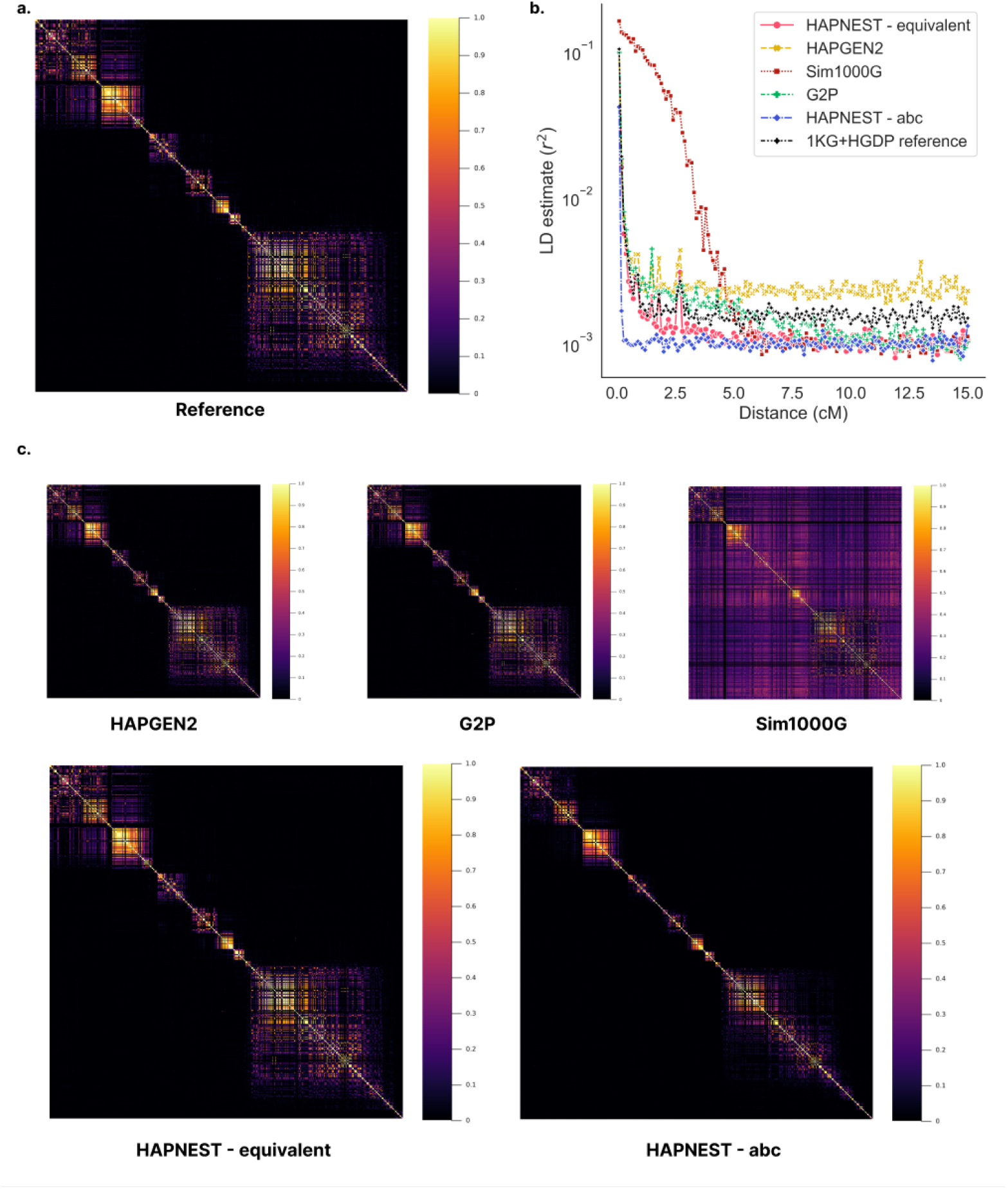
**a**. LD correlation for 500 contiguous SNPs selected at random from chromosome 21 HapMap3 variants, for the European-ancestry reference dataset (*N_ref_* = 775); **b**. Comparison of LD decay for *N_syn_* = 1000 European-ancestry synthetic samples; **c**. Comparison of LD correlation (for same 500 SNPs shown in reference panel) for *N_syn_* = 1000 European-ancestry synthetic samples

**Figure 4:**
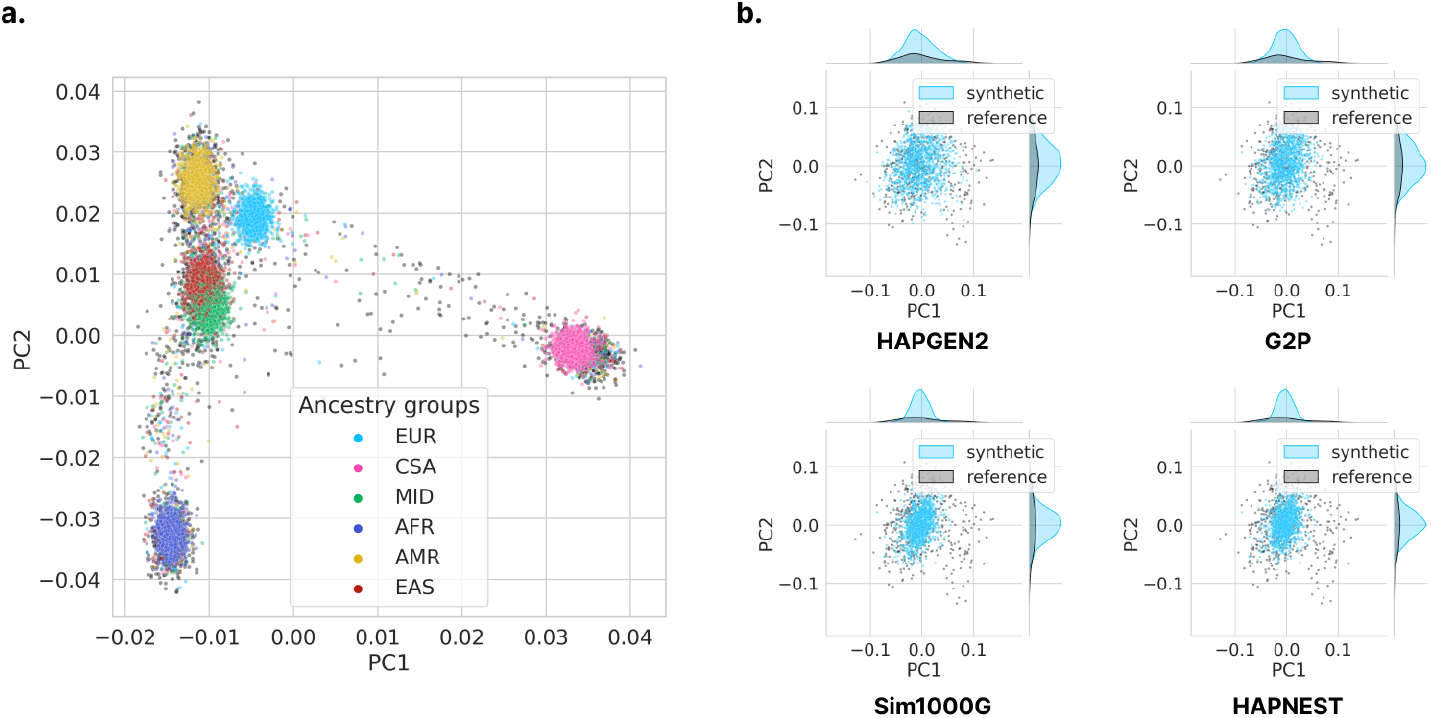
**a**. PCA projection plot for *N_syn_* = 10000 synthetic samples generated by the HAPNEST method, for chromosome 21 HapMap3 variants; **b**. Comparison of PCA projection plots and bivariate densities for *N_syn_* = 1000 European-ancestry synthetic samples. The highest PC alignment score for preservation of population structure is 0.311 for HAPGEN2, followed by 0.243 (HAPNEST-equivalent), 0.222 (G2P), 0.182 (HAPNEST-abc) and 0.043 (Sim1000G)

#### 2.3.2 Generalisability and diversity

HAPNEST-abc reached the best generalisability and diversity of all methods evaluated (Supplementary, Table 2, 3) when considering *N_syn_* = 1000 synthetic samples. However, it is more appropriate to measure generalisability and diversity on larger and more realistic sample sizes. As there is a limited number of haplotypes in the reference dataset, one might expect that when generating thousands of synthetic samples, some generated genomes might eventually be copies of or highly related with genomes in the reference set. As shown in the next section, scalability is an issue for Sim1000G and G2P, so in this experiment we only consider HAPNEST and HAPGEN2. To evaluate the impact of the size of the reference panel, we consider both the full reference (*N* = 775) and a smaller reference (*N* = 100). We observe that HAPNEST outperforms HAPGEN2 for both generalisability and diversity on larger sample samples (Figure 5). These results are not a function of the synthetic data sample size for either method, due to simplifying assumptions of the statistical models used by these methods. However, the generalisability and diversity performance is affected by the size of the reference data. We also demonstrate that for both methods, generalisability and diversity can be improved by increasing the number of reference samples.

**Figure 5:**
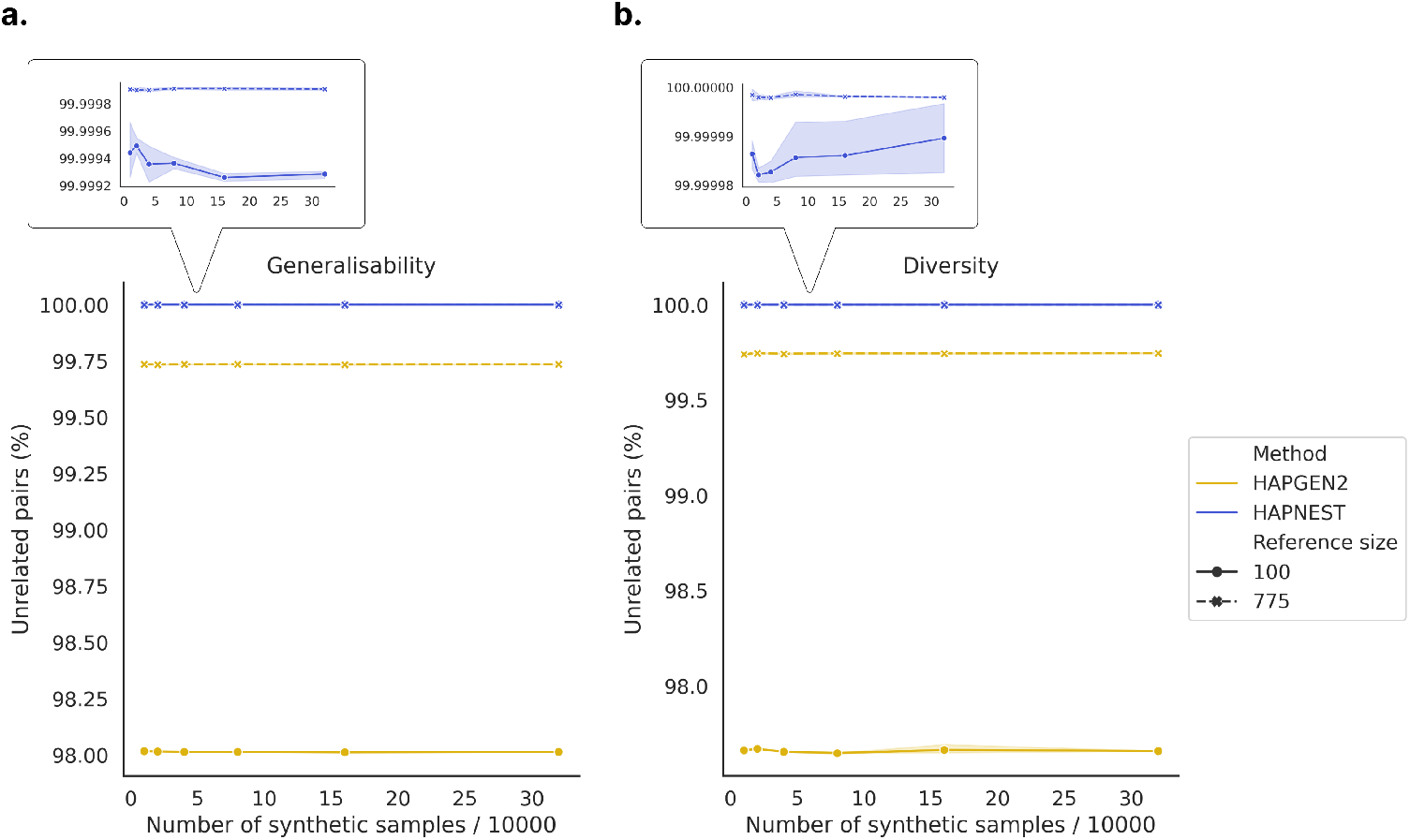
The **a**. generalisability and **b**. diversity scores for two reference sizes (*N_ref_* = 100 and *N_ref_* = 775) and various sample sizes, averaged across five trials for chromosome 21 HapMap3 variants, and the HAPNEST and HAPGEN2 methods. The ratio *N/N_e_* is fixed to ensure a fair comparison with the same average segment lengths. Generalisability is calculated as 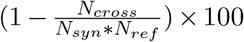, where *N_cross_* is the number of closely related pairs (i.e. twins or first-degree relatives, as determined by the kinship coefficient) between the reference and synthetic datasets. Diversity is calculated as 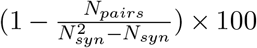, where *N_pairs_* is the number of closely related pairs in the synthetic dataset.

### 2.4 Scalability analysis for large sample sizes

The scalability of HAPNEST is validated by measuring the computational speed of generating genotype datasets for a range of sample sizes, compared with the widely used HAPGEN2 software tool. The other two methods (Sim1000G and G2P) are excluded from this comparison as they did not scale to the large sample sizes considered here. We observe that while generation times are similar for small sample sizes, HAPNEST is increasingly faster than HAPGEN2 for larger sample sizes (Figure 6) which approach the size of modern biobank-scale genetic datasets. This gain in computational speed is achieved by a more efficient algorithm and its efficient multi-threaded implementation in the Julia programming language.

**Figure 6:**
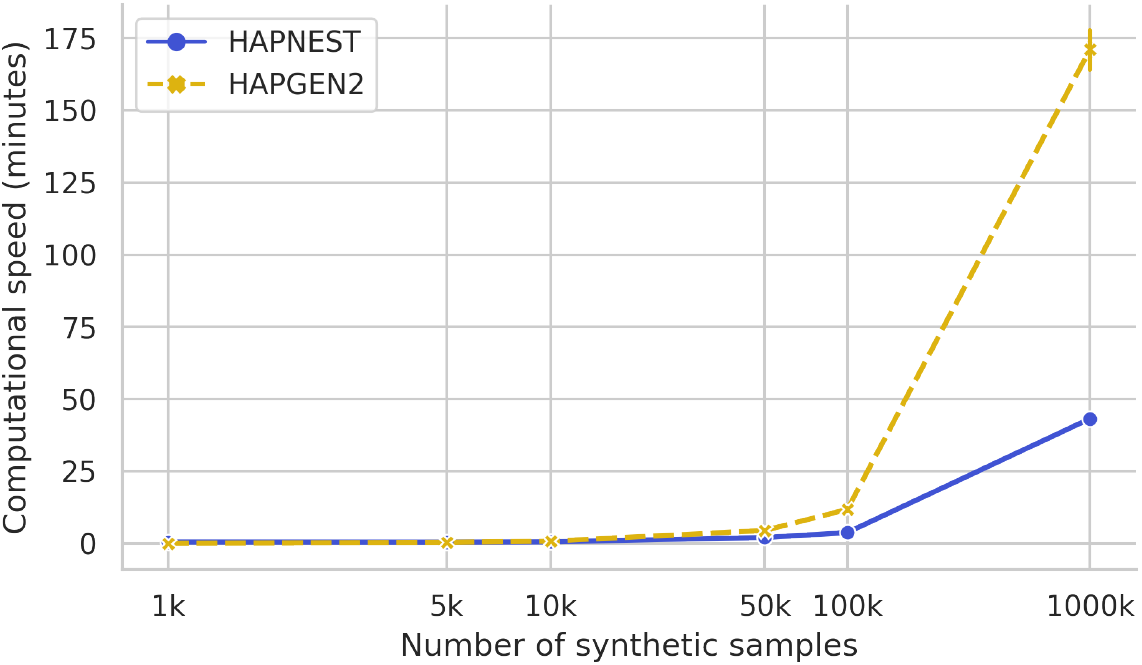
Simulation times for genotype datasets for HAPNEST and HAPGEN2 (other methods are excluded from this comparison due to scalability and compatibility issues), averaged for five trials with error bars plotted, for chromosome 21 HapMap3 variants. The comparison was performed on Intel Xeon Gold 6230 2.1 GHz processors with 8 cores and 32GB RAM. Since the simulation time depends on the input configuration, the experiment is controlled by setting *ρ* to the average recombination rate used by HAPGEN2 (*ρ* = 2.185 for chromosome 21), and using *N_e_* = 500 for both methods (to eliminate bias from mutations).

### 2.5 Overview of phenotype generation methods

A continuous or binary phenotype can be assigned to each sample as an aggregation of genetic effect, user-input covariate effect (if any) and environmental noise. The genetic component is generated as a weighted sum of causal allele counts (Figure 1, Panel d). For each causal SNP *β_i_*, the effect size is drawn from a Gaussian distribution with 0 mean and variance determined by three well-studied factors impacting heritability of the variants, the minor allele frequency (MAF) *p_i_*, local linkage structural *r_i_*, and the functional annotation *s_i_* of the SNP:

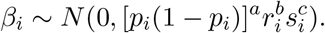

Power parameters *a*, *b*, and *c* reflect strength of negative selection on each aspect and we used extensive empirical observations [20, 21, 22] to chose the default parameters. HAPNEST allows SNP’s effect sizes to be drawn from a mixture of distributions with different width, corresponding to variable level of heritability. Our model also allows flexible assignment of individual components’ contribution to the phenotype (heritability), as well as the number of causal variants constituting the genetic risk (polygenicity). We run GWASs for 50,000 synthetic individuals and 1,049,096 HapMap 3 SNPs based on phenotypes generated under different genetic architectures. The Manhattan plots visually resemble Manhattan plots obtained on real data with similar heritability and polygenicity (Figures 7, 13 and 14). Figure 7 shows exemplary GWAS results for traits under two extreme scenarios: low heritability, low polygenicity, and high heritability, high polygenicity. The former resembles phenotypes such as atrial fibrillation and flutter (Figure 13), and the latter resembles typically more heterogeneous traits, such as body pain (Figure 14). Our approach allows us to specify genetic correlations between phenotypes within and, importantly, between ancestry groups.

**Figure 7:**
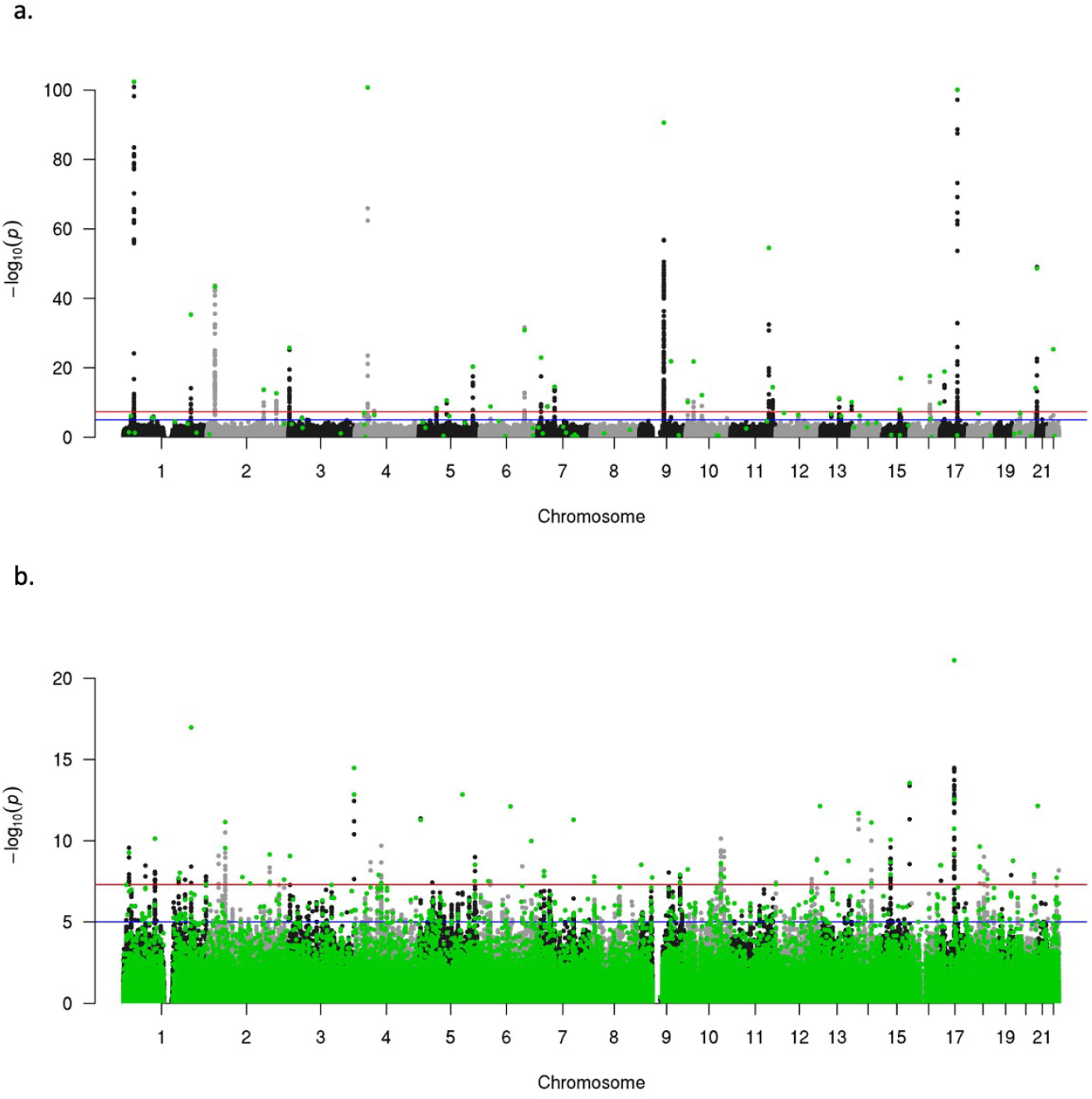
Example GWAS Manhattan plots for phenotypes under various genetic architectures. Colored in green are causal SNPs on trait liability under each setup. a. Phenotype with low heritability (0.1) and low polygenicity (0.0001, i.e. approximately 0.01% of total SNPs having causal effects on trait liability); b. Phenotype with high heritability (0.9) and high polygenicity (0.1, i.e. approximately 10% of total SNPs having causal effects on trait liability).

### 2.6 Application: Comparison of polygenic risk scoring methods

We demonstrated the utility of HAPNEST by comparing 7 PRS methods using synthetic data from 5 ancestry groups. We first generated a synthetic training dataset of 100,000 individuals of European ancestries, and performed a standard GWAS using software plink2 [23], correcting for top 20 PCs. We subsequently used the summary statistics to build PRSs in a separate synthetic test set of 25,000 individuals (5,000 samples from each ancestry group). To demonstrate variability across genetic architectures, GWAS summary statistics are computed for nine continuous phenotypic traits, with varying heritability (0.03, 0.1, 0.5) and polygenicity (0.0001, 0.005, 0.1). We assumed a genetic correlation of 1 across all ancestry groups.

The evaluation of the PRS methods is based on the reference-standardised framework of Pain et al. [4], where for continuous traits, the PRS performance is measured in terms of Pearson correlation between the predicted and observed values. The optimal parameters for each PRS method are identified using cross validation (CV), or pseudovalidation (PseudoVal), if CV is not available.

Better predictive performance is observed for higher heritability, lower polygenicity architectures (Supplementary, Figure 15). No single PRS method was observed to perform best across all genetic architectures. Methods with sparsity-inducing shrinkage priors (e.g. PRScs) were observed to perform better for higher heritability, lower polygencity architectures, where genetic effects on most SNPs are zero (Figure 8, Panel c), while other approaches such as MegaPRS performed better for lower heritability, higher polygenicity architectures (Figure 8, Panel a). Multi-ancestry results replicate known issues with transferability of polygenic risk scores based on European-ancestry summary statistics (Figure 8, Panels b and d).

**Figure 8:**
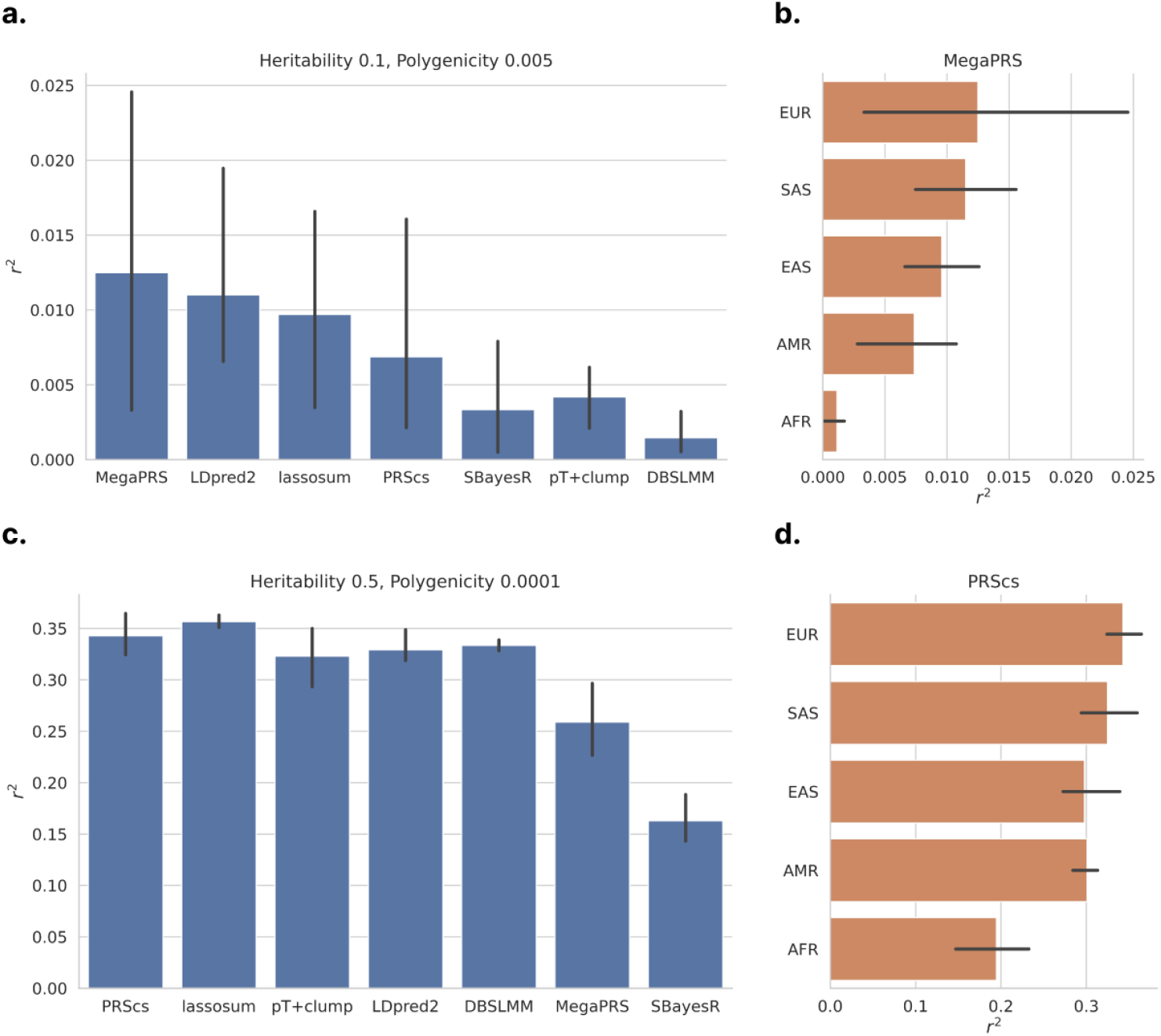
PRS results for two genetic architectures, averaged across 3 experiment trials with error bars showing the range of outcomes, for HapMap3 variants across 22 chromosomes. **a**. Pearson correlation between predicted and observed values, for various PRS methods and a European-ancestry phenotype with heritability 0.1 and polygenicity 0.005. **b**. Pearson correlation for various target ancestry groups for the best-performing PRS method (MegaPRS) for the heritability 0.1 and polygenicity 0.005 phenotype. **c**. Pearson correlation between predicted and observed values, for various PRS methods and a European-ancestry phenotype with heritability 0.5 and polygenicity 0.0001. **d**. Pearson correlation for various target ancestry groups for the best-performing PRS method (PRScs) for the heritability 0.5 and polygenicity 0.0001 phenotype.

## 3 Discussion

In this study, we proposed HAPNEST, a new algorithm to generate realistic individual-level genetic and phenotypic data and provide an efficient implementation. HAPNEST meets the demand for diverse, biobank-scale genomic data by improving scalability compared to existing methods. Users can customise population parameters or use parameter estimates derived from the reference dataset. Previous studies have been inconsistent in their approach to evaluating the quality of the generated synthetic data. We provide a comprehensive set of measures to be used for data quality evaluation that have been proposed in the statistical genetics and differential-privacy literature [24]. Genotype generation, phenotype generation and evaluation modules are wrapped in user-friendly Docker or Singularity containers, where each module can be run independently.

Synthetic genotypes are generated by copying and assembling haplotype segments from the reference genome, with distribution of segment length determined by specifics of the target population, including recombination rates, effective population size and samples in the reference panel. Parameters are optimised through the ABC algorithm, which typically results in an output dataset well-balanced across fidelity and generalisability metrics. On top of that, we introduced mutations to the synthetic genome to reduce similarity across individuals. Our approach is, in spirit, similar to HAPGEN2, but to improve computational scalability and generalisability we have introduced modeling of varying, rather than constant, coalescence time, and the use of mutation ages to determine if mutations are present in synthetic samples.

From our systematic evaluations and experiments, we noticed some general trade-offs in synthetic data quality and in the parameter selection. One trade-off occurs between the preservation of population LD structure and synthetic sample relatedness when constructing large synthetic datasets from much smaller reference datasets. Our observations indicated that parameters optimising the preservation of LD usually result in higher levels of sample relatedness, as LD typically comes with larger average segment length copied from the reference. On the other hand, shortened segments allow more combinations and higher sample level variability, which results in samples that are less related to each other but increased fragmentation in the LD structure. Furthermore, smaller segments lead to more computational input/output operations when constructing synthetic data files and a slight increase in running time. Segments copied from the reference genome in our algorithm can be conceptually viewed as identity-by-descent (IBD) segments in population genetics [25, 26]. As can be seen in equation 2, recombination events (*ρ_s_*) happen over time (*T*) in the population. Thus, IBD segments degrade over time, which also shows an impact on LD [27, 28]. Our algorithm also provides an implementation of generating “admixed” samples by sampling from multiple reference populations under user defined compositions. However, we would like to note that this approach does not accurately reflect the process of multi-population diverging and intermixing, therefore it should be used and interpreted carefully.

Compared to other methods, HAPNEST-generated genotypes demonstrated better diversity and generalisability which are essential features when scaling to large sample sizes. While the genetic relatedness analysis indicated that the genotypes are sufficiently different from the reference data, a nearest-neighbour adversarial accuracy close to 0.5 indicates that statistically speaking, it would be difficult to discern a synthetic genotype from a real genotype. These properties of synthetic datasets are desirable in the context of data privacy, where we may want to create a synthetic twin of sensitive data that preserves key statistical properties of the real data, but cannot be traced back to real individuals. However, we note that the criteria used in our analysis are not sufficient for differential privacy guarantees, and we advise to use HAPNEST, or any of the reference-based generation methods, only on publicly-available genomics datasets.

Once individual level genotypes have been generated, we can subsequently assign phenotypes to each sample as an aggregation of polygenic effects, non-genetic effects and environmental noise. We also implemented population-specific phenotypic effects by assuming shared causal variants across populations with distinct but correlated effect sizes, and multi-trait simulation allowing for different genetic correlation and pleiotropy models.

We believe our tool can benefit the community especially for GWAS related method development, for which one of the examples can be PRS computation and evaluation. HAPNEST allows researchers to assess the validity of genetic scoring methods under a broad variety of setups, including cross-ancestry, trans-diagnostic, and different genetic architectures. Here, as a demonstration of its utility, we applied PRSpipe^2^ to synthetic data generated by HAPNEST and found that our results, to a great degree, replicated what has been observed by Pain et. al [4]. As widely discussed, we found lower cross-ancestry portability of PRSs derived in a single ancestry. For a given phenotype, we set genetic correlations between ancestry groups to 1 and this might be higher than what is observed in real settings and result in slightly inflated trans-ethnic PRS prediction performance. Nevertheless, we still observed reduced prediction accuracy in non-European samples, indicating the synthetic genotype captured the differences of MAF and LD structures across populations. Results under different genetic architectures are concordant with the general expectation: we observe better performance of PRS for phenotypes with higher heritability and lower polygenicity due to the existence of few variants with larger effect that explain large amounts of phenotypic variance. We also noticed that the best performing method can depend on different genetic architecture, reflecting the need for careful considerations when choosing a PRS method. As more studies come online that examine the clinical utility of PRSs, it will be important to have a reference dataset where old and new PRS methods can be compared and their robustness can be assessed as a function of the genetic and phenotypic architecture. We used HAPNEST to create one of the largest genomics synthetic datasets today including 1 million individuals across 6 major continental ancestry groups, 6.8 million variants and 9 phenotypes. We hope this dataset can generate a reference set for deriving and testing PRS methods within a unified framework.

## 4 Data availability

We have made available a pre-simulated synthetic dataset for 1,008,000 individuals and 9 continuous phenotypic traits for over 6.8 million SNPs and 6 ancestry groups at https://www.ebi.ac.uk/biostudies/studies/S-BSST936. There is also a smaller example dataset available at this link.

## 5 Code availability

The HAPNEST software is available at https://github.com/intervene-EU-H2020/synthetic_data. The software can be used to simulate synthetic datasets and evaluate synthetic data quality.

## 6 Acknowledgements

This project has received funding from the European Union’s Horizon 2020 research and innovation programme under grant agreement No 101016775.

## Footnotes

2 PRSpipe is a Snakemake pipeline developed to calculate and evaluate polygenic risk scores from GWAS summary statistics. It implements and extends the GenoPred [4] pipeline, a reference standardized framework for the prediction of PRS using various state-of-the-art methods.

