## Supplementary material for "HAPNEST: efficient, large-scale generation and evaluation of synthetic datasets for genotypes and phenotypes"

### A Supplementary methods

We present a comprehensive software pipeline for synthetic data generation, evaluation and optimisation, which for ease of portability and reproducibility, has been made available as Docker and Singularity containers (Supplementary, Figure 9). The data generation workflow implements efficient methods for simulating large-scale genotype and phenotype datasets. The evaluation workflow computes a set of quantitative metrics and visualisations for assessing the quality of synthetic datasets. Finally, the optimisation workflow can be used to examine the posterior distributions of model parameters, to aid with model selection.

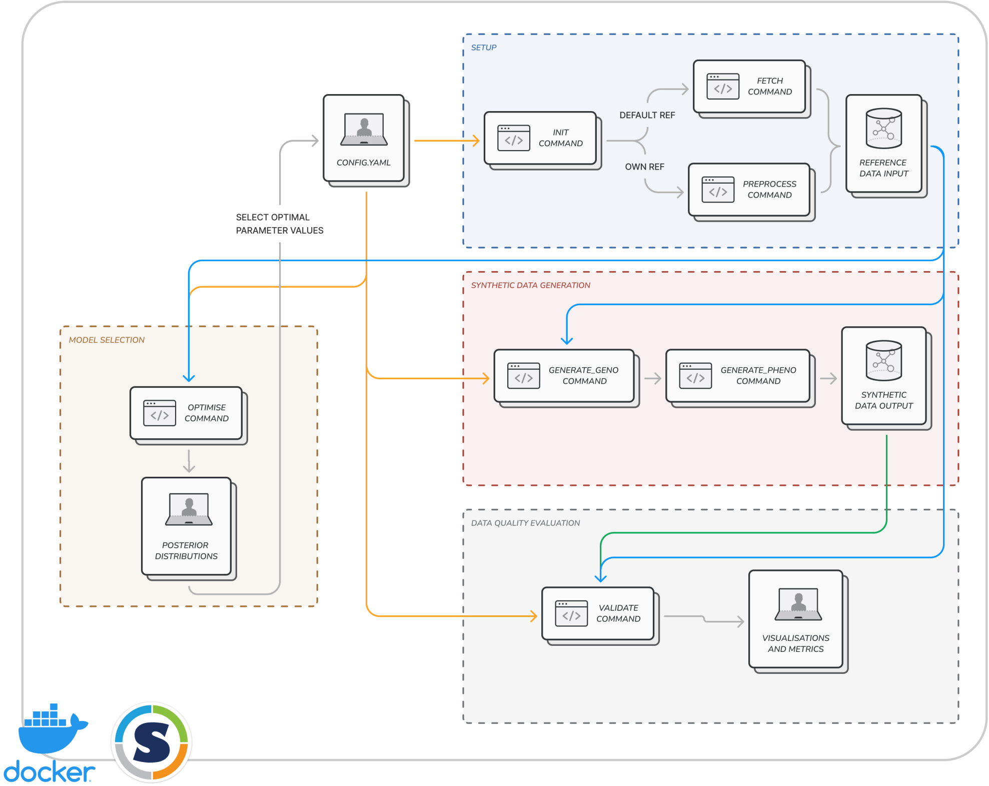


Figure 9: Architecture diagram for the software tool, available at `https://github.com/intervene-EU-H2020/synthetic_data`. Given a YAML configuration file and reference dataset as input, the software tool provides modules for preprocessing the reference data, generating synthetic datasets, evaluating data quality, and selecting optimal model parameters. The collection of modules has been containerised using both Docker and Singularity for ease of use.

#### A.1 Genotype generation

##### A.1.1 Algorithm overview

A synthetic genotype, $g$, is generated as a pair of synthetic paternal and maternal haplotypes, where each synthetic haplotype, $h_i$, $i \in \{1, 2\}$, is constructed as a mosaic of segments of various lengths imperfectly copied from real haplotypes. The real haplotypes to copy from are sampled uniformly from a reference dataset, $\mathcal{D}_s$, limited to individuals belonging to a certain ancestry group $s \in S$. The ancestry groups $S$ should be interpreted in the context of the original reference datasets from which they were derived [3].

Alternatively, users can specify the proportion of real haplotypes to sample from each ancestry group, but we refer to the Discussion section of the paper regarding the complications in interpreting admixed samples. Segments of length $\ell$ (in centimorgans) are sampled from the real haplotypes (Figure 1, Panel b) based on a simplified stochastic model of the coalescent and recombination processes,

$$\ell \sim Exp(2T\rho_s),\ T \sim Gamma(2, N_s/N_{e,s}), \tag{2}$$

where $\rho_s$ is the population-specific recombination rate, $N_{e,s}$ is the population-specific mean effective population size, and $N_s$ is the number of reference samples for population $s$. The simulation of a coalescence time, $T$, is one aspect in which HAPNEST differs from previous methods such as HAPGEN2. Another feature we introduce is that to reduce close copying of genotypes from the reference, the presence of a genetic variant at position $i$ is only copied if $T \leq m_i$, where $m_i$ is the variant's age of mutation (obtained from [16]). Two synthetic haplotypes, $h_i$, $i \in \{1, 2\}$, constructed in this way are added element-wise to create a synthetic genotype, $g$ (Figure 1, Panel c).

The algorithm is implemented efficiently in the Julia programming language, using a reference dataset of 4062 genotypes from the publicly available 1000 Genomes Project and Human Genome Diversity Project datasets. This data is downloaded from the Genome Aggregation Database (gnomAD) [17] and filtered to retain the HapMap3 variants. Users of the HAPNEST software can choose to download this pre-processed reference or process their own using a module included as part

[3] $S$ = {AFR, AMR, EAS, EUR, CSA, MID}, where AFR=African , AMR=Admixed American, EAS=East Asian, EUR=European, CSA=Central/South Asian, MID=Middle Eastern

of the HAPNEST software. To enable biobank-scale data generation, the number of computers, threads of execution, and memory constraints can be configured so that HAPNEST utilises the available computing resources. This enables highly parallelisable, multi-threaded generation of millions of samples for millions of variants in a matter of hours.

##### A.1.2 Posterior distributions of model parameters

We use approximate Bayesian computation (ABC) rejection sampling to estimate the posterior distributions of the unknown model parameters, $\rho_s$ and $N_{e,s}$, for each $s \in S$. ABC involves simulating many synthetic datasets and compares the real and synthetic data sets using a discrepancy between the two datasets, giving a high probability to parameter values that yield a small discrepancy. This discrepancy is measured by $d(Q(\mathcal{D}_{real}), Q(\mathcal{D}_{synthetic}))$, where $d$ is Euclidean distance and $Q$ is a vector concatenating LD decay and cross-relatedness (the proportion of first degree and second degree relatives, between the real and synthetic datasets for $\mathcal{D}_{synthetic}$ and between two random partitions of the real dataset for $\mathcal{D}_{real}$).

This approach favors models that preserve LD structure between the real and synthetic datasets, while limiting the degree of cross-relatedness to that found within the reference data to encourage diversification and generalisation of the synthetic samples. For computationally expensive simulations of large synthetic datasets, our software pipeline runs ABC with a small number of simulations by training a Gaussian process regression model to estimate the remaining simulation results [29].

#### A.2 Phenotype generation

**Overview** Once the genotype has been generated, we subsequently assign liability of phenotypes for each individual as a summation of genetic effect, effect of user input covariates, and noise $\epsilon$:

$$Y = G + U + \epsilon,$$

where $Y$ is liability of the phenotype being simulated, $G$ is total genetic effect and $U$ is total effect of all covariates. Users can specify proportion of variance for each component in phenotype construction. Further, based on the given trait prevalence, binary case-control statues can be assigned as below:

$$Y_b = \begin{cases} 1 & \text{if } Y > \Phi^{-1}(1-k) \\ 0 & \text{otherwise} \end{cases},$$

where $Y_b$ is the Bernoulli random variable denoting samples case/control statues, $k$ is the trait population prevalence and $\Phi$ is the cumulative density function of standardised normal distribution.

**Genetic liability of single trait** The genetic liability of an individual for a given trait is defined as weighted sum of risk allele counts, where the weights are causal effect sizes randomly sampled from a normal distribution with zero mean. Users can choose to input a list of causal SNPs, or customise the polygenicity parameter for the phenotype simulation, which is defined as the proportion of total SNPs being causal (ie. Polygenicity = 0.01 indicates approximately 1% of total SNPs will have a causal effect on the trait determination). Since heritability of a SNP can be affected by many properties, such as SNP's minor allele frequency (MAF) or local linkage structure, we model the variance of effect size distribution for any causal SNP as a function of its MAF, LD score, and functional annotation

$$\beta_i \sim N(0, [p_i(1-p_i)]^a r_i^b s_i^c),$$

where $\beta_i$ is the effect size for causal SNP $i$, $p_i$ is MAF of the SNP, $r_i$ is its local LD score, and $s_i$ is its functional annotation index. Due to the computation intensity from calculating local LD score for all causal SNPs from the input genome, especially when the polygenicity is high, by default our tool uses precomputed LD score from African population in Hapmap 3 project [9] as a reference, which is subject to user customisation. Adjusting parameters $a$, $b$ and $c$ changes the relationship between SNP heritability and those properties, which in reality, is an indication of negative selection strength. As an example, choosing $a = 0$ will make the SNP effect size distribution invariant to its MAF, and same with $b$ for LD score and $c$ for SNP annotation. On top of these, we also allow stratifying the causal SNP effects into multiple levels (e.g. separating SNPs with large, medium, small effects) by adopting a Gaussian mixture distribution with components having different distribution width as below

$$\beta_i \sim \sum_{j=1}^{k} \pi_j N(0, \sigma_j^2 [p_i(1-p_i)]^a r_i^b s_i^c), \ \sum_{j=1}^{k} \pi_j = 1,$$

where $k$ is the number of components and $\sigma_j^2$ is parameter controlling the distribution width for the $j-$th component, both can be specified by the user. Each SNP falls into one of the mixture components $j$ with probability $\pi_j$, which is determined by the reciprocal of the width parameter $\sigma_j^2$ and scaled so that we have $\sum_{j=1}^{k} \pi_j = 1$. For a more realistic setup, we suggest using a wide range of width parameters, covering small amount of SNPs each having relatively higher heritability, to massive SNPs with each harbouring lower heritability [30].

**Genetic effects on different populations** Our phenotype generation also encompasses population specific genetic effects, not only through incorporating MAF differences across population, but also by introducing the cross-ancestry genetic correlation parameter. We adopt the common assumption that for the same phenotype, causal variants are shared across ancestry but not necessarily their exact effect sizes, and generate correlated base causal effects for different populations from multivariate normal distribution. On top of the base effect size, we further integrate effect of the population specific MAF of the SNP. To be more specific, for each causal SNPs, we first generate population based effect sizes $\beta_{\mathbf{i}}$ from a multivariate Gaussian mixture distribution

$$\beta_{\mathbf{i}} \sim \sum_{j=1}^{k} \pi_j MVN(\mathbf{0}, \sigma_j^2 \mathbf{\Sigma})$$

where $\mathbf{\Sigma}$ is the population covariance matrix with each element $[\mathbf{\Sigma}]_{mn}$ denoting covariance of standardised SNP effects in population $m$ and $n$. As mentioned in the previous section, these base effect sizes can also be stratified in to various levels, using the component width parameter $\sigma_j^2$. Then, the final effect size of SNP $i$ in a population $m$ will be

$$\beta_{im}^{*} = \beta_{im} \times \sqrt{p_{im}([1-p_{im})]^a r_i^b s_i^c}$$

where $p_{im}$ is the allele frequency of SNP $i$ in population $m$.

**Simulating multiple correlated phenotypes** Our tool also allows generation of multiple phenotypes with predefined genetic correlation and pleiotropic model simultaneously. Each trait has its own heritability and polygenicity. Moreover, SNP effects on different traits can be correlated under user-defined coefficient. Unlike the cross-ancestry genetic correlation, different traits can have different set of causal SNPs. In this case, the pleiotropy parameter can be specified to determine the proportion of causal SNPs being shared across different traits. Note that the phenotype correlation parameter applies to correlation of pleiotropy SNP effects, as well as effects of non-genetic covariates and environmental noise.

#### A.3 Evaluation

To evaluate the quality of data generated by the synthetic data generation algorithm, we consider three key aspects of data for downstream tasks.

1. Fidelity: Realistic synthetic genetic data should retain the key summary statistics of the original genetic data.

2. Diversity: Synthetic genetic data should be sufficiently different from the real-world genetic data, to preserve privacy.

3. Validity: The synthetic phenotypes should behave similarly to the real-world phenotypes in down-stream genetics applications.

For each of these aspects, we design and evaluate multiple metrics to analyse the reliability as well as the robustness of the synthetic data generation algorithm.

##### A.3.1 Fidelity of synthetic samples

We compare four properties of the synthetic dataset against real dataset to understand the fidelity of synthetic samples in retaining key statistical properties.

1. **Minor Allele Frequency** We measure the discrepancy of Minor Allele Frequency (MAF) from the original dataset using two statistical distances. First, we consider the observed value of each haplotype as an independent Bernoulli random variable. Hence, a genomic sample can be seen as a collection of realised Bernoulli random variables with different parameters.

Considering real data as one realisation and synthetic data as another realisation of this collection of random variables, we can compute statistical discrepancies between these two random variables. We use KL divergence and Wasserstein distance between Bernoulli random variables as the metric to measure the discrepancy in MAF.

2. **Population structure** Most of the downstream PRS tasks are interested in analysing the effect of SNPs at a population level, it is extremely critical to maintaining the genetic structure of real population in the synthetic samples. We use the popular concept of Principal Component Analysis to ensure that the synthetic samples are well aligned with the real samples. In order to quantify the misalignment between the population structures, we designed a new metric 'PC alignment'. Let $\{\boldsymbol{x}_i\}_{i=1}^{P}$ be the set of principal components in real data, where $\boldsymbol{x}_i \in \mathbb{R}^{n_{SNP}}$ and $\{\tilde{\boldsymbol{x}}_i\}_{i=1}^{P}$ be the principal components of synthetic dataset. We can measure alignment between two principal components using the cosine distance defined as $d_i = \frac{\boldsymbol{x}_i^T \tilde{\boldsymbol{x}}_i}{|\boldsymbol{x}||\tilde{\boldsymbol{x}}_i|}$. If the $i^{th}$ principal components are well aligned, $|d_i| \to 1$, else $|d_i| \to 0$. Hence, cosine distance gives us a tool to quantify the degree to which the principal components are aligned. Further, we weigh the cosine distance between principal components by the proportion of variance explained by that PC in real data, denoted by $\eta_i$. Hence, the final value is calculated using the formula,

$$\zeta = \sum_{i=1}^{K} \eta_i \frac{\boldsymbol{x}_i^T \tilde{\boldsymbol{x}}_i}{|\boldsymbol{x}||\tilde{\boldsymbol{x}}_i|}, \tag{3}$$

where top $K$ components are considered for computational simplicity.

3. **Linkage Disequilibrium** While measuring MAF preserves the probability of occurrence on each SNP, the co-occurrence of neighbouring SNPs are not ensured due to the independent and identically distributed (IID) assumption. In order to verify the local structure of co-occurrence is maintained in the synthetic dataset, we compute the Linkage Disequilibrium correlation matrix and compare them between real data and synthetic data.

4. **Nearest-Neighbour Adversarial Accuracy** A sufficiently good synthetic data sample must be indistinguishable from the real data sample. Based on this principle, we use the adversarial accuracy [24] of a classifier to quantify the closeness of synthetic dataset to real

dataset. Let $N$ be the number of real samples as well as artificial samples, $\mathbb{1}(\cdot)$ evaluates to 1 if the condition is true, else 0, $d_{TS}(i)$ is the distance between $i^{th}$ real datapoint and its nearest neighbour in synthetic dataset, $d_{ST}(i)$ is the distance between $i^{th}$ synthetic datapoint and its nearest neighbour in real dataset, $d_{TT}(i)$ is the distance to nearest neighbour of $i^{th}$ real data point in the real dataset and $d_{SS}(i)$ same as above in synthetic dataset, then Nearest Neighbour Adversarial Accuracy $AA_{TS}$ defined as,

$$AA_{truth} = \frac{1}{n}\sum_{i=1}^{n} \mathbb{1}(d_{TS}(i) > d_{TT}(i)) \tag{4}$$

$$AA_{syn} = \frac{1}{n}\sum_{i=1}^{n} \mathbb{1}(d_{ST}(i) > d_{SS}(i)) \tag{5}$$

$$AA_{TS} = \frac{1}{2}\left(AA_{truth} + AA_{syn}\right) \tag{6}$$

Here, $AA_{truth}$ measure the fraction of real samples for which nearest neighbour in metric space is a synthetic sample and $AA_{syn}$ measure the same fraction for synthetic samples. When the synthetic data is very close to real data, we will not be able to distinguish real data samples from synthetic samples and hence $AA_{TS} \to 0.5$.

###### A.3.2 Diversity of synthetic samples

A main aim of creating synthetic datasets is to preserve the privacy of individual samples in the original dataset, yet allowing the scientific community to develop new tools from the vast amount of data available. We study the 'closeness' of synthetic samples to real samples using two key metrics.

1. **Kinship analysis**: For kinship analysis [31], we compute the kinship between samples within the generated dataset as well as kinship across the reference dataset and generated dataset. Within dataset kinship analysis is useful in analysing the diversity in the synthetic samples within the cohort and kinship across datasets are useful to address the privacy concerns of reference data leaking into synthetic data as well as quantifying the between synthetic and real samples.

2. **IBS analysis** Identical-by-state (IBS) analysis shows samples which share identical sequence at a particular locus and is useful in analysing the closeness between samples. We analyse

the distribution of IBS values to compare the real data with generated synthetic data.

##### A.3.3 Validity of phenotypes

To ensure the phenotype generation algorithm returns realistic phenotype for each individual given the input parameters, we created phenotypes for 5,000 European ancestry samples under a range of heritability 0.1, 0.3, 0.5, 0.7, 0.9 and polygenicity 0.0001 or 0.1. Assuming a prevalence of 0.1, for each continuous phenotype, we also assigned binary case/control labels for all individuals. We computed heritability using genomic-relatedness-based restricted maximum-likelihood method implemented in GCTA (CGTA-GREML) [32] and compared the estimated heritability to our input parameters. Under each condition, the genetic relationship matrices (GRM) used for heritability estimation was computed using causal SNPs underlying the simulated phenotypes. We also calculated the genetic correlation of two traits simulated under various pleiotropic relationships using LDSC [22] as another benchmark of the phenotype generation.

#### B Supplementary experiments

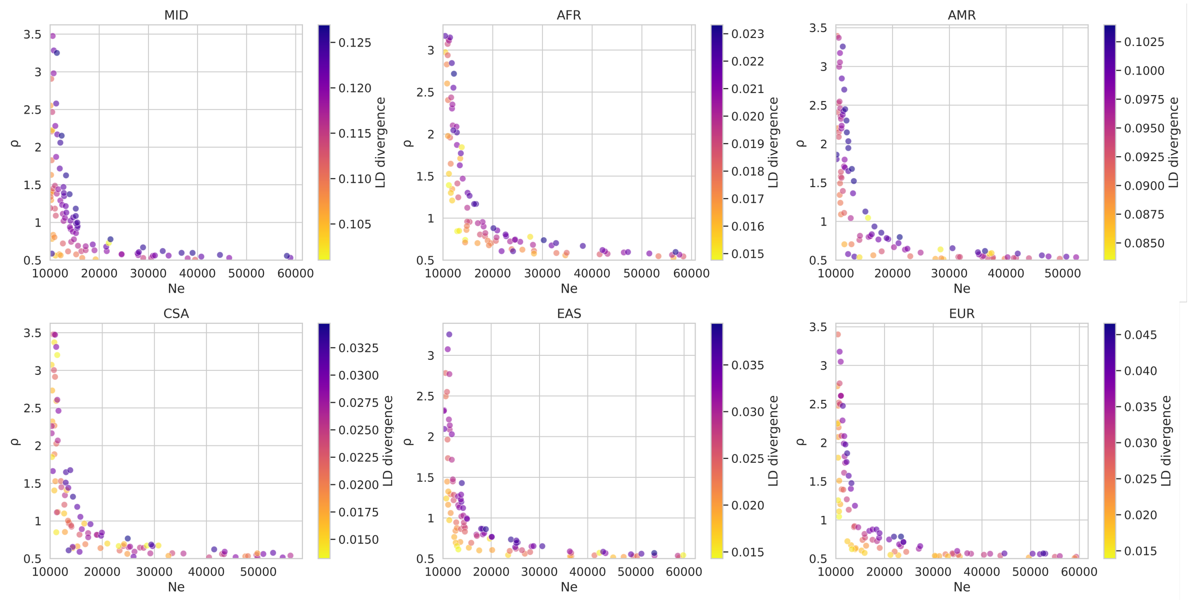


Figure 10: Scatter plots corresponding to the posterior distribution samples plotted in Figure 2. The colormap represents the Euclidean distance between the LD decay curves of the simulated datasets and the real dataset (fidelity objective).

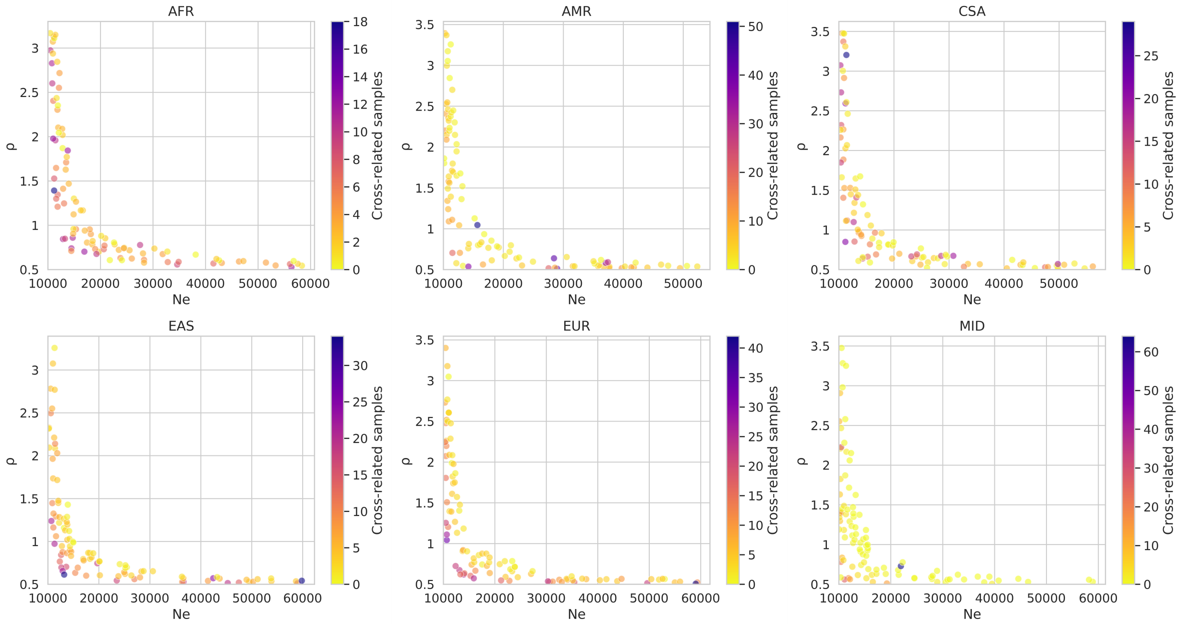


Figure 11: Scatter plots corresponding to the posterior distribution samples plotted in Figure 2. The colormap represents the number of samples in the simulated datasets that have close (twin or first degree) relatives in the real dataset (generalisability objective).

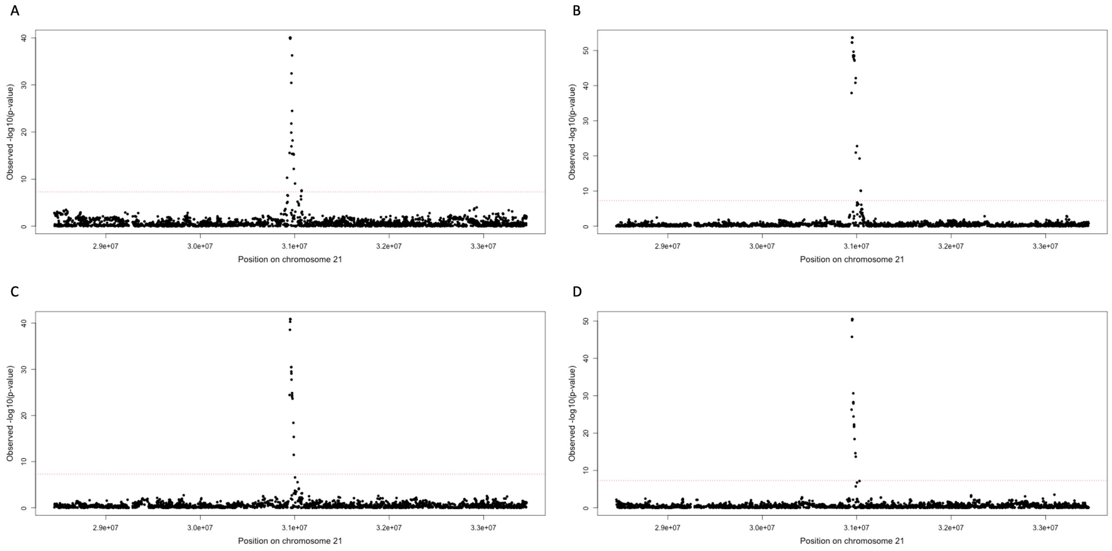


Figure 12: **Regional plot around causal SNP on chromosome 21 under different genotype generation parameters, comparing with real reference genome.** GWAS ran on different genomes with phenotype generated under same parameters and randomization seed: heritability = 0.05 and single causal variant on chromosome 21. Each GWAS group contains 4062 EUR samples, which is the sample size for the reference genome of same ancestry. Panel **A.** shows the regional plot for GWAS ran on the real reference genome. The rest three panels each shows a representative case when different aspects are prioritized during parameter selection. **B.** Genotype simulated with $rho = 0.35, Ne = 8000$, where fidelity is prioritized; **C.** Genotype simulated with $rho = 0.68, Ne = 11700$, where fidelity and generalisability are balanced; **D.** Genotype simulated with $rho = 0.75, Ne = 25000$, where generalisability is prioritized. See table 6 for quantified comparison of GWAS hits.

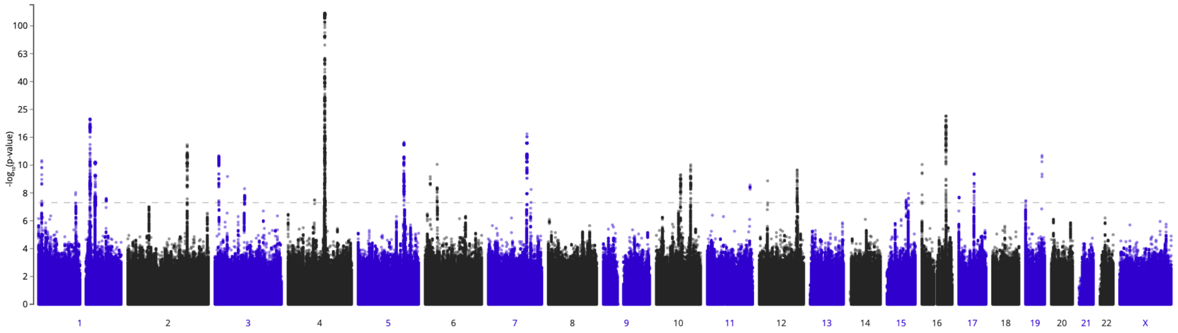


Figure 13: Manhattan plot of atrial fibrillation and flutter from Finngen freeze 5. GWAS included 22,068 cases and 116,926 controls.

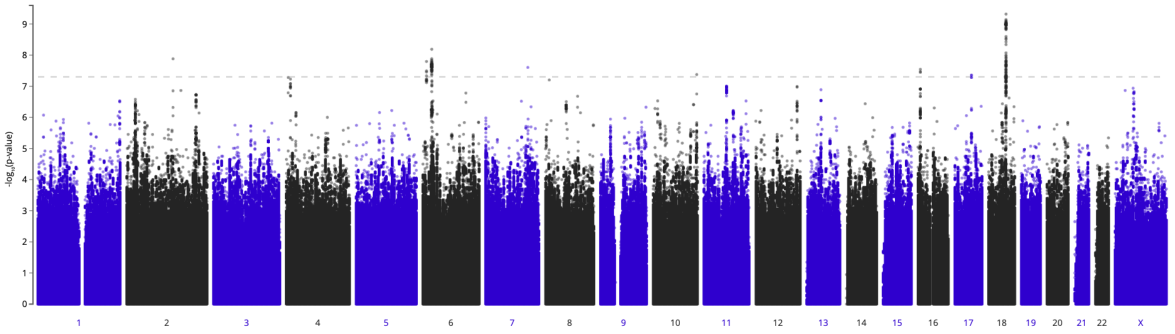


Figure 14: Manhattan plot of pain (limb, back, neck, head abdominally) from Finngen freeze 5. GWAS included 87,242 cases and 131,127 controls.

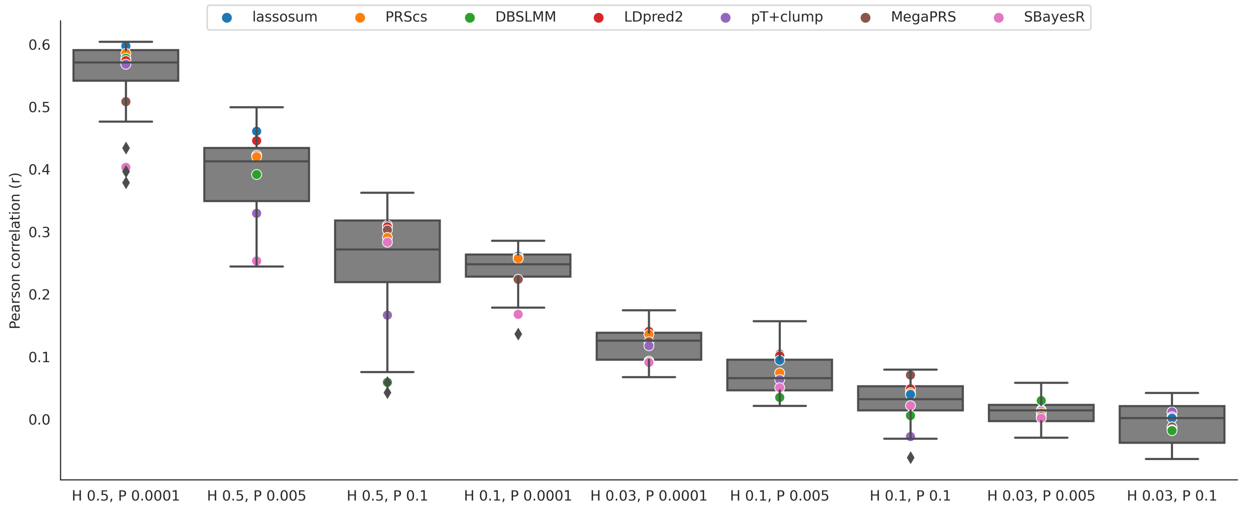


Figure 15: Pearson correlation between predicted and observed values, for various PRS methods and European-ancestry phenotypes with various genetic architectures (H corresponds to the heritability value and P corresponds to the polygenicity value). The optimal parameters for each PRS method are identified using cross validation (CV), or pseudovalidation (PseudoVal), if CV is not available. Scatter plots show the average value obtained for each PRS method, from 3 iterations of the experiment, for HapMap3 variants across 22 chromosomes.

| | **HAPGEN2** | **G2P** | **Sim1000G** | **HAPNEST abc** | **HAPNEST eq.** |
|---|---|---|---|---|---|
| **NN adversarial accuracy** | 0.623 | **0.505** | 0.706 | 0.559 | 0.333 |
| **LD decay** | 0.014 | 0.013 | 0.442 | 0.066 | **0.012** |
| **MAF** | 0.020 | **0.013** | 0.014 | 0.014 | **0.013** |
| **PC alignment** | **0.311** | 0.222 | 0.043 | 0.182 | 0.243 |

Table 1: Comparison of the evaluation metrics nearest neighbor adversarial accuracy, LD decay (Euclidean distance between LD decay vectors), MAF (Wasserstein divergence) and PC alignment, for four synthetic data generation tools. Each synthetic dataset contains 1000 samples, based on a European ancestry reference panel for HapMap3 variants on chromosome 21. HAPNEST-abc uses parameter values determined by the ABC algorithm for balancing the LD and relatedness objectives, while HAPNEST-equivalent uses parameters that are more equivalent to those used for the other methods ($\rho_s = 0.05, N_{e,s} = 2000$). Best results are indicated in bold.

| | **Dataset relatedness** | | | | |
|---|---|---|---|---|---|
| | **HAPGEN2** | **G2P** | **Sim1000G** | **HAPNEST abc** | **HAPNEST eq.** |
| **MZ twins** | 2 | **0** | **0** | **0** | **0** |
| **1st degree** | 2600 | 3 | 2492 | **0** | 1 |
| **2nd degree** | 381 | 916 | 42277 | **0** | 188 |
| **3rd degree** | 16904 | 11332 | 52144 | **4785** | 16163 |
| **Unrelated** | 479613 | 487249 | 402587 | **494715** | 483148 |

Table 2: Comparison of relatedness metrics for four synthetic data generation tools, measuring the number of genetically related genotype pairs within the synthetic datasets. Best results are indicated in bold.

| | Cross relatedness | | | | |
|---|---|---|---|---|---|
| | **HAPGEN2** | **G2P** | **Sim1000G** | **HAPNEST abc** | **HAPNEST eq.** |
| **MZ twins** | 1 | **0** | **0** | **0** | **0** |
| **1st degree** | 2031 | 7 | **0** | **0** | 106 |
| **2nd degree** | 672 | 1234 | 4982 | **21** | 709 |
| **3rd degree** | 18832 | 17281 | 58898 | **11068** | 21056 |
| **Unrelated** | 731464 | 734478 | 689120 | **741911** | 731129 |

Table 3: Comparison of cross-relatedness metrics for four synthetic data generation tools, measuring the number of genetically related genotype pairs between the synthetic and reference datasets. Best results are indicated in bold.

| Simulation input | | | LDSC genetic correlation | |
|---|---|---|---|---|
| Heritability | Polygenicity | Trait correlation | with causal SNPs | without causal SNPs |
| 0.3 | 0.0001 | 0.1 | 0.2344 (0.25) | 0.2546 (0.23) |
| 0.3 | 0.0001 | 0.5 | 0.4847 (0.20) | 0.4662 (0.20) |
| 0.3 | 0.0001 | 0.9 | 0.9472 (0.19) | 0.9459 (0.11) |
| 0.3 | 0.1 | 0.1 | 0.1122 (0.06) | 0.0926 (0.057) |
| 0.3 | 0.1 | 0.5 | 0.5161 (0.04) | 0.5147 (0.04) |
| 0.3 | 0.1 | 0.9 | 0.8943 (0.01) | 0.8956 (0.01) |
| 0.7 | 0.0001 | 0.1 | -0.3132 (0.19) | -0.3198 (0.18) |
| 0.7 | 0.0001 | 0.5 | 0.3478 (0.15) | 0.3355 (0.14) |
| 0.7 | 0.0001 | 0.9 | 0.8621 (0.08) | 0.8545 (0.08) |
| 0.7 | 0.1 | 0.1 | 0.1291 (0.032) | 0.1260 (0.029) |
| 0.7 | 0.1 | 0.5 | 0.5179 (0.024) | 0.5165 (0.02) |
| 0.7 | 0.1 | 0.9 | 0.8989 (0.01) | 0.8969 (0.01) |

Table 4: **Genetic correlation of simulated traits with and without causal variants.**

| | polygenicity = 0.0001 | | | polygenicity = 0.1 | | |
|---|---|---|---|---|---|---|
| Input heritability | GCTA estimate for continuous trait | GCTA estimate for binary trait observed | liability | GCTA estimate for continuous trait | GCTA estimate for binary trait observed | liability |
| 0.1 | 0.0975 (0.01) | 0.0304 (0.01) | 0.0912 (0.02) | 0.1202 (0.05) | 0.05 (0.05) | 0.1362 (0.14) |
| 0.3 | 0.2890 (0.03) | 0.1012 (0.02) | 0.3010 (0.05) | 0.2692 (0.05) | 0.12 (0.05) | 0.3507 (0.14) |
| 0.5 | 0.5063 (0.04) | 0.1629 (0.02) | 0.4926 (0.07) | 0.5313 (0.05) | 0.12 (0.05) | 0.3441 (0.14) |
| 0.7 | 0.7055 (0.03) | 0.2375 (0.03) | 0.6784 (0.07) | 0.6016 (0.05) | 0.21 (0.05) | 0.6186 (0.14) |
| 0.9 | 0.8984 (0.01) | 0.3168 (0.03) | 0.9375 (0.09) | 0.8662 (0.04) | 0.31 (0.05) | 0.8928 (0.14) |

Table 5: **Input and CGTA estimated heritability for synthetic data with different polygenicity.**

| $p$-value threshold | $\mathbf{5 \times 10^{-8}}$ | $\mathbf{5 \times 10^{-7}}$ | $\mathbf{5 \times 10^{-6}}$ | $\mathbf{5 \times 10^{-5}}$ |
|---|---|---|---|---|
| **Real reference genome** | 22 | 25 | 27 | 38 |
| $\mathbf{rho = 0.35, N_e = 8000}$ | 22 | 25 | 28 | 37 |
| $\mathbf{rho = 0.68, N_e = 11700}$ | 18 | 19 | 21 | 22 |
| $\mathbf{rho = 1.2, N_e = 25000}$ | 17 | 19 | 21 | 22 |

Table 6: **Comparison of SNPs surpassing various $p$-value thresholds in GWAS for genomes generated under different parameters.** Also see figure 12. Reporting number of SNPs with GWAS $p$-value lower than each threshold.
